# Allostery between distant structural regions dictates selectivity in GPCR:G protein coupling

**DOI:** 10.1101/2025.08.04.668482

**Authors:** Elizaveta Mukhaleva, Edgardo J. Sánchez Rivas, Sergio Branciamore, Andrei S. Rodin, Sivaraj Sivaramakrishnan, Nagarajan Vaidehi

## Abstract

Selectivity in GPCR-G protein coupling is widely regarded as a solved problem that derives from a combination of structural hotspot pairings and dynamics at the protein-protein interface. Here, using interpretable machine learning Bayesian Network model with Molecular Dynamics simulations and experiments, we reveal the influence of distant residue communities within the G protein core on coupling selectivity. We observed distinct cooperative hotspot residues across different Gα protein subtypes, including key regions such as the N-terminus, h4s6 loop, and H5 helix. These results demonstrate the intricate allosteric dependencies between the core and the H5 helix in stabilizing selective interactions. The functional significance of these cooperative regions is validated through subtype swapping mutations. By introducing targeted Gαq like mutations in the Gαs core, we successfully altered receptor coupling profile to signal through Gαq. Our findings emphasize that cooperative interactions in the Gα core are not only crucial for selectivity but can also be leveraged to engineer G proteins with tailored coupling preferences.

**Teaser:** Allosteric interactions in the G protein core shape GPCR selectivity and can be engineered to rewire receptor signaling preferences.

## Introduction

Upon agonist binding, G-protein coupled receptors (GPCRs) couple to trimeric G proteins that trigger cellular signaling pathways (1). GPCRs form the largest superfamily of membrane proteins, mediating responses to diverse stimuli, and represent a major class of drug targets (1– 4). The binding of agonists to GPCRs activates the receptor to couple to specific Gα protein subtypes such as Gαs, Gαq, Gαi, or Gα12/13. This Gα subtype-specific interaction shapes the downstream signaling outcomes, thereby influencing functional specificity in various physiological and pathological contexts (3). The factors that affect the selective coupling of GPCRs to G proteins are multi-fold including structural features as well as cellular factors.

Analysis of the high-resolution structures of GPCR:Gα protein complexes offers a glimpse of the interacting residues in the interface but is not sufficient to decipher selectivity determinants since the interface is dynamic (4–7). This is evident from alanine scanning mutation studies combined with G protein coupling measurements showing that the residues that do not make direct contact in the GPCR:Gα protein interface in the three-dimensional structures still significantly affect the receptor activity upon mutation (3, 4, 6). Understanding how GPCRs selectively couple to specific Gα proteins despite overlapping interface features is a critical step toward designing therapeutics that can precisely modulate signaling pathways.

Multiple studies including ours have shown that the dynamic properties of protein interfaces, including GPCR:Gα protein interactions, are critical for Gα protein coupling specificity (7–13). The last 10 amino acid residues in the H5 helix are known as the “tip” and the rest of the entire Gα protein is referred to as the “core” (4, 14). Many studies have highlighted the importance of the “tip” in Gα protein coupling selectivity. Other studies have shown that the “core” (rest of the Gα protein without the tip) of the Gα protein is as important in coupling selectivity (4) as the tip. However, the specific residues and/or structural regions in the core of the Gα protein that confer selectivity are not known. Additionally, there is a serious lack of understanding of how different structural regions within the core of the Gα protein allosterically affect the GPCR:Gα protein coupling interface (15, 16).

In this study, we have used an interpretable machine learning method, Bayesian Network (BN) Modeling, to perform a fully data-driven analysis of the extensive molecular dynamics (MD) trajectories of GPCR:Gα protein contacts combined with experiments to uncover the specific structural regions and residues in the core of the Gα protein that confer G protein coupling selectivity to GPCRs. Our results highlight the previously unknown allosteric cooperativity between the H5 helix, N-terminus and h4s6-loop in the core of the Gα subunit.

## Results

In our previous study, we developed the BN modeling method to analyze the all-atom MD simulation trajectories in a fully data-driven way to uncover the cooperative residue pair interactions in the GPCR:Gα protein interface (13). We define “cooperative” interactions as the residue interactions in the GPCR:Gα protein interface that show probabilistic co-dependencies with other interaction residue pairs in the interface. Such cooperative interactions, when mutated, will lead to a cascading effect on the dissociation of G proteins from GPCRs.

The BN modeling is an interpretable unsupervised machine learning method that offers a promising solution for analyzing cooperative interactions in dynamic systems. Details of the BN models as applied to MD simulation trajectories have been published (17). Briefly, given a set of input variables, the BN model recovers a directed acyclic graph representing the joint probability distribution of these variables, depicted as nodes in the graph (Fig. S1A). Unlike traditional network analyses, which rely on spatial proximity or correlation metrics (18–25), BN modeling identifies direct dependencies, filtering out spurious correlations (17). This makes it particularly suitable for studying coordinated or collective behavior of GPCR:Gα protein interface interactions or the cooperative dynamics at protein-protein interfaces. In this study, the residue pairs in the GPCR:Gα interface are represented as nodes (random variables) and their direct dependencies as directed edges. Thus, the recovered BN model summarizes the co-dependencies among the GPCR:Gα residue interactions. The workflow used in this work for generating the BN models of GPCR:Gα protein interactions is shown in Fig. 1.

**Figure 1.**
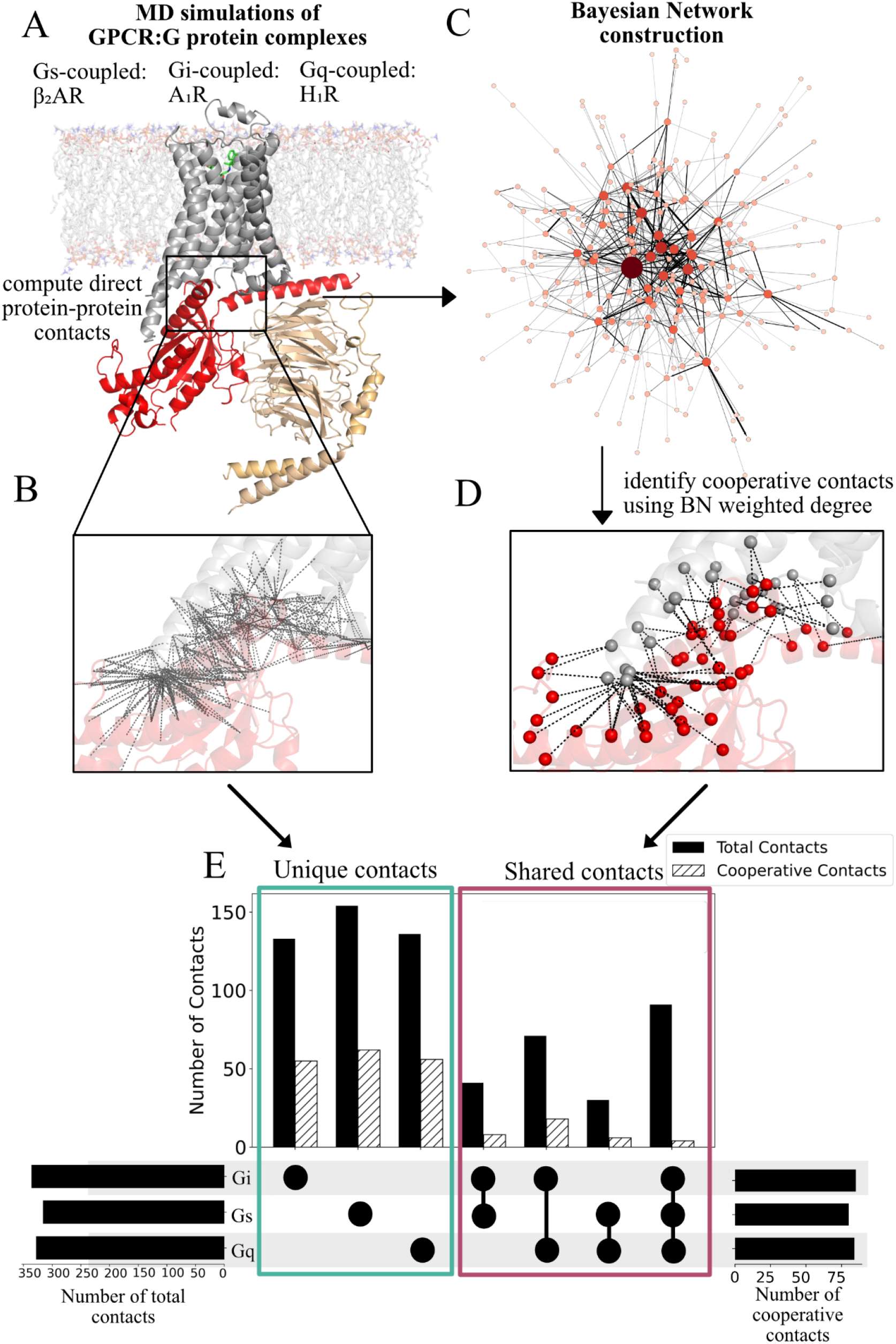
Bayesian Network Analysis shows shared and G protein-specific cooperative residue contacts in the GPCR:Gα interface. **A.** MD simulations of GPCR:Gα protein complexes performed in POPC bilayer were used to obtain GPCR:Gα interface residue contacts matrixes using the program “GetContacts” (32). **B**. The GPCR:Gα protein residue contacts in the interface were calculated in every MD snapshot and used as input to the BN modeling software BaNDyT. **C**. Bayesian Network model of GPCR:Gα interface contacts showing the probabilistic co-dependencies among these contacts. **D**. The GPCR:Gα protein interface residue contacts that show a high weighted degree in the BN model. **E**. The upset plot of the number of high-ranked (top quartile) GPCR:Gα protein interface contacts that are common among the six GPCRs studied here and are distinct for each Gα protein subtype.

Starting from their respective three-dimensional structures, we performed all-atom MD simulations, spanning a total of 5μs, in explicit POPC bilayer for each of the six GPCR:Gα protein complexes: 2 Gαs coupled receptors, namely β_2_-adrenergic receptor (β_2_AR) (pdb id: 3SN6 (26)) and adenosine 2A receptor (A_2A_R) (6GDG (27)), 2 Gαq coupled receptors, histamine receptor (H_1_R), angiotensin II receptor 1 (AT_1_R) (H_1_R: 7DFL (28), AT_1_R: 7F6G (29)), and 2 Gαi coupled receptors, Adenosine 1 receptor (A_1_R) and serotonin 1B receptor (5HT_1B_R). (A_1_R: 6D9H (30), 5HT_1B_R: 6G79 (31)) (Fig. 1A, Fig. S1A). The details of the MD simulations are given in the Methods section. For each GPCR:Gα protein complex, we generated pairwise contact information for the residues in the GPCR:Gα protein interface using GetContacts (32) software, resulting in a superset of 300 to 400 contacts in the interface for all the six GPCR complexes (Fig. 1B). Each contact fingerprint matrix represents GPCR:Gα residue contacts in binary form (1 or 0, contact present or absent in each MD snapshot). These binary matrices were inputs to our MD-centric BN modeling software, BaNDyT (33). For each GPCR:Gα protein complex, a BN probabilistic graphical model was recovered (Fig. 1C). In the BN model shown in Fig. 1C, each node represents a pair of interacting residues in the GPCR:Gα protein interface and the edges connecting the nodes show the direct probabilistic dependencies between the nodes. We calculated the sum of all the edge strengths of each node, the weighted degree, which is a measure of the co-dependencies of the other nodes on this node (Fig.1D). The nodes with a high weighted degree can be conceptualized as cooperative contacts since they show high dependencies with other nodes (13, 17).

Comparison of the total residue contacts in the GPCR:Gα protein interface and that of the cooperative contacts among Gαs, Gαq and Gαi (top 25 percentile of the contacts by the weighted degree score) for the six GPCR:Gα protein systems is shown in Fig. 1E. More than 50% of the GPCR:Gα protein contacts are shared among the Gαs, Gαi, and Gαq coupling systems. The percentage of shared contacts is 60.5% for Gαi coupled receptors, 58.5% for Gαq coupled receptors, and 51.2% for Gαs coupled receptors. Interestingly, the cooperative contacts are largely specific to Gα subtype. 64.7% of cooperative contacts are from Gαi specific contacts, 66.7% in the Gαq specific contacts, and 88.6% are from Gαs specific contacts. These findings suggest that Gα specific interactions drive cooperativity in the GPCR:Gα protein interface, which may contribute to the selectivity of GPCR:Gα protein coupling.

### How do Cooperative Contacts Affect G Protein Selective Coupling to GPCRs?

The total number of cooperative contacts in various structural regions of the GPCRs and Gα proteins are shown in the heat map in Figures 2A-C. The high cooperativity contacts are between H5 helix of the G protein and the TM5 and TM6 residues in the GPCRs for both Gαi and Gαq coupled complexes. In the Gαi coupled complexes, hns1, and s2s3 loops (Fig. 2A) contacts with intracellular loop 2 (ICL2) contain cooperative contacts. The cooperative contacts are predominantly located in the H5 helix in Gαi, in the H5 helix, hns1 loop, and h4s6 loops for Gαq coupled complexes, and in the H5 helix, hgh4 region, H4 helix, and h4s6 loop for Gαs coupled complexes (Fig. 2C). Taken together, these analyses highlight that H5 helix contains cooperative contacts that are common among Gα subtypes. The s2s3 loop is prominent in Gαi but absent in Gαq, while the hgh4 region contains cooperative interactions in Gαs but not in Gαq or Gαi coupled complexes This differential pattern of cooperativity reinforces the hypothesis that these regions could be contributing to the selective interaction between the receptor and Gα proteins, allowing for selectivity in coupling.

**Figure 2.**
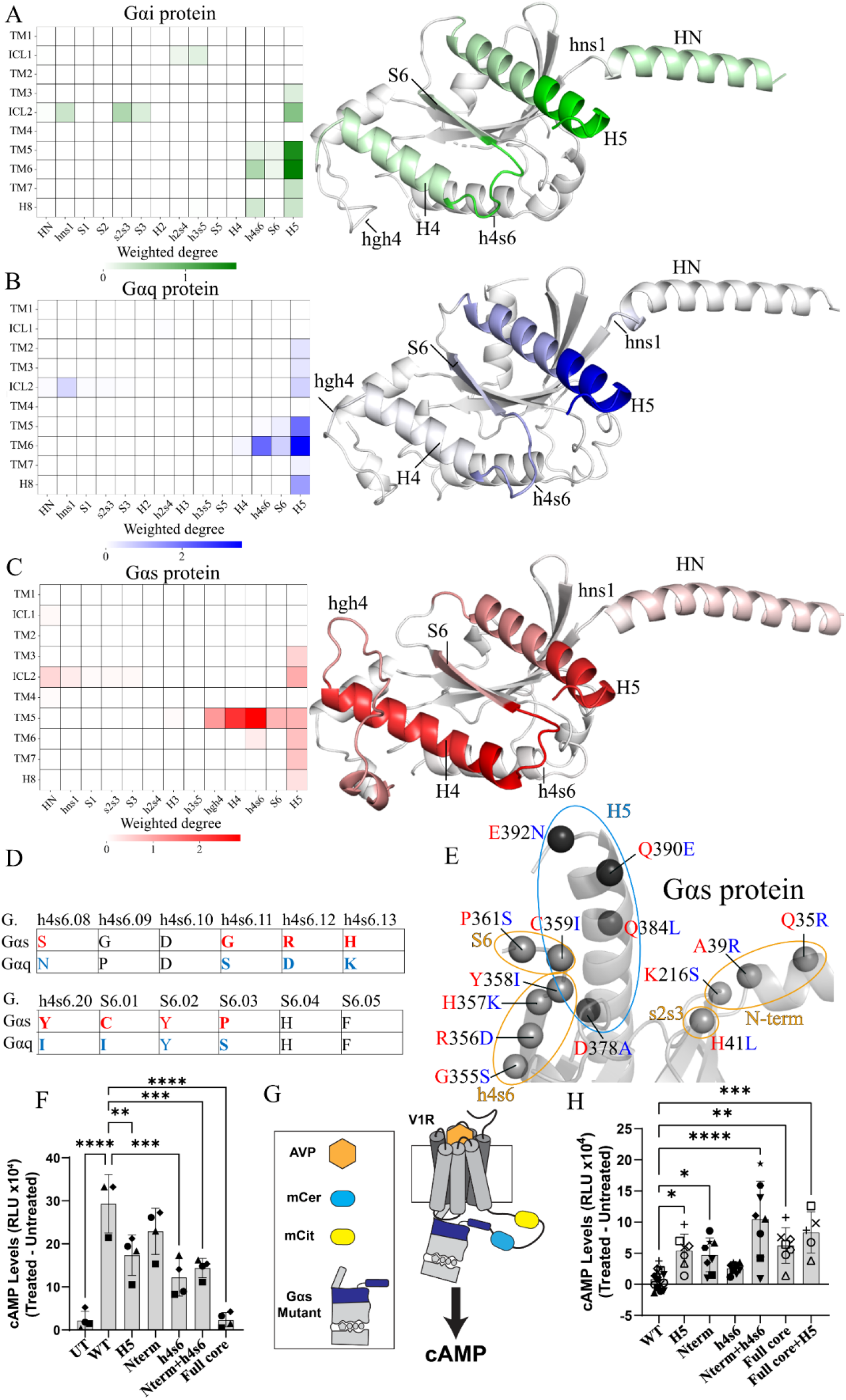
Selectivity Switching in GPCR-G Protein Complexes. **A–C.** Distribution of cooperative contact regions across different Gα protein subtypes. **A**. Heat map of the weighted degree (cooperativity) of the GPCR:Gα interface contacts in different GPCR and Gα protein structural regions for Gαi coupled complexes (left). The structure of Gαi protein with cooperativity regions mapped in green (right); intensity of color is proportional to the weighted degree. **B**. Cooperativity heat map for Gαq coupled complexes (left), and structure of Gαq protein with cooperativity regions mapped in blue (right). **C**. Cooperativity heat map for Gαs coupled complexes (left), the structure of Gαs protein with cooperativity regions mapped in red (right). **D**. Section of the multiple sequence alignment of h4s6 and S6 structural elements of Gαs and Gαq proteins. Colored residues (Gαs – red, Gαq – blue) indicate the residue positions with high cooperativity (top 25% of contacts ranked by weighted degree). Non-conserved residues between Gαs and Gαq are highlighted in bold. **E**. Structural model of the Gαs protein showing predicted mutation positions, with core residues colored in light grey and H5 helix residues highlighted in dark grey. Individual segments listed in Table 1 are circled. **F**. Secondary messenger cAMP production in HEK293ΔGsix cells showing that Cterm, Nterm, h4s6, Nterm+h4s6, and full core mutants lose coupling to β_2_AR. Significance is denoted as **P < 0.01, ***P < 0.001, ****P < 0.0001. **G**. Schematic of SPASM sensor assembled with mCerulean, allowing expression measurements of the construct, and a flexible linker, allowing movement and function of the G protein. **H**. Secondary messenger cAMP production in HEK293 cells showing that Cterm, Nterm, h4s6, Nterm/h4s6, and full core mutants gain coupling to Vasopressin receptor, V_1_R. Significance is denoted as *p < 0.05, **p < 0.01, ***p < 0.001, ****p < 0.0001.

As mentioned before, the H5 helix of the Gα subunit is a well-established structural determinant of GPCR-G protein coupling selectivity, stabilizing the GPCR-G protein complex (4, 7, 11). However, the Gα protein core, encompassing residues beyond the last 10 amino acids, also plays a critical role in selectivity through regions such as the αN-β1 hinge and switch I/II, which mediate receptor-specific and subfamily-specific interactions (4, 34). Despite these insights, the individual residues within the core contributing to selective coupling remain undetermined. Additionally, chimeras made using tips of different Gα subtypes with a different core are assumed to behave as the subtype of the tip thus neglecting the allosteric influence of the different structural regions in the core on the tip. To identify how specific cooperative structural regions might contribute to Gα protein coupling selectivity, we designed a strategy aimed at switching the coupling profile from Gαs to Gαq coupled receptors. We selected Gα residues to be mutated using the following criteria: (i) The residues with high cooperativity score (weighted degree) in both Gαs and Gαq and (ii) these residues must be different in the amino acid type in both these proteins as shown in the sequence alignment in Fig. 2G and Fig. S2A. These criteria ensured that we targeted residues critical for cooperative interactions while maximizing the chances of switching coupling selectivity. Based on these criteria, we have predicted the residue positions shown in Table 1 for Gαs to Gαq switching mutations in the core of the Gα protein and tested them experimentally as described below.

**Table 1.**
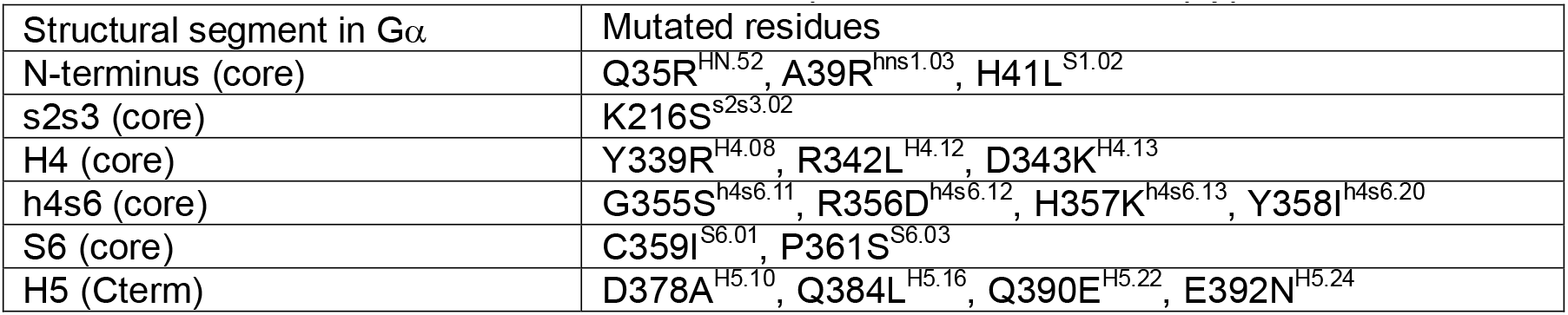
Predicted Mutants in the core of the Gαs protein to switch to Gαq type.

To address the functional integrity of the mutations of the whole set of core residues in Gαs (all the mutants in the N-terminus, s2s3, H4, h4s6, and S6 – Table 1) to Gαq type, a mCerulean-tagged Gα protein mutant was expressed in HEK293ΔGsix, HEK293 cell absent of six functional Gα proteins (Gs, Golf, Gq/11, G12/13). We observed a significant decrease in the expression of the mutant compared to wild-type mCerulean-tagged Gαs (Fig. S2B). Additionally, this mutant did not accumulate cyclic adenosine monophosphate (cAMP) when the cells were exposed to forskolin with no significant difference to non-transfected HEK293ΔGsix (Fig. S2C) showing inactivity. To identify which specific region mutation(s) reduced the expression and contributed to the depletion of the forskolin signal, we tested the mutants in individual structural regions shown in Table 1. We observed that recovering H4 segment mutations in Gαs showed no statistical difference in expression and cAMP accumulation compared to the wild-type Gαs (Fig. S2C). Since the mutations in the H4 helix led to non-functional Gαs protein we eliminated these from further experiments.

We tested the ability of Gαq-like Gαs mutants to accumulate cAMP upon isoproterenol (iso)-induced activation of HEK293ΔGsix endogenous β2-adrenergic receptor (β2AR). As shown in Fig. 2E, the overexpression of H5 helix Gαq-like mutants made in Gαs was able to significantly reduce isoproterenol-induced cAMP accumulation. Furthermore, the Gαq-like full-core mutant disrupts the iso-induced cAMP, resembling the non-transfected condition (UT). This result suggests that cooperative interactions within both the core and H5 helix regions are essential for efficient signaling through Gαs. Moreover, the loss of function observed in the full-core mutant supports the notion that these structural segments in the core contribute cooperatively to maintaining the structural integrity necessary for optimal Gαs signaling.

To further explore whether these mutations could confer Gq-like signaling properties, we tested each mutant’s ability to couple to a Gq-coupled receptor, the vasopressin 1A receptor (V_1A_R). We tether the G protein to a V_1A_R linked to a SPASM sensor, allowing high local concentration and one-to-one stoichiometry between receptor and G protein (5). As illustrated in Fig. 2G, the SPASM sensor is assembled with mCerulean, allowing expression measurements of the construct, and a flexible linker, allowing movement and function of the G protein. As V_1A_R is primarily a Gq-coupled receptor, ligand-induced cAMP accumulation is not expected. Therefore, the transition from Gαs-to-Gαq on H5 and core segments will drive the coupling of functional Gαs into a Gαq-coupled receptor. The results in Fig. 2H indicate an increase of AVP-induced V_1A_R signaling through Gαs for H5, full-core, and H5+core containing mutants. This gain of function suggests that residues involved in the cooperativity between β_2_AR and Gαs are also involved in the interface between V_1A_R and Gαq.

In summary, our findings reveal that the cooperativity landscape across GPCR:Gα protein complexes is distinct for each Gα subtype, with specific cooperative regions that are unique to each Gα protein. By engineering mutations based on these cooperativity hotspots, we successfully modulated the coupling profile of Gαs, highlighting the potential of cooperative regions to influence selectivity. These results provide a framework for understanding the structural basis of Gα protein selectivity and suggest that targeting cooperative sites could be a promising strategy for designing Gα protein variants with altered coupling preferences.

### Allosteric regulation of the H5 helix by different structural regions in the core of the Gα protein

BN models provide information on the allosteric co-dependencies between different structural regions of the protein. To quantify and test such allosteric regulation of the tip of the Gα protein by different structural regions in the core, we calculated the weighted degree for each structural region of the Gα protein encompassing all nodes containing H5-helix contacts, which are highlighted in blue in Figure 3A.

**Figure 3.**
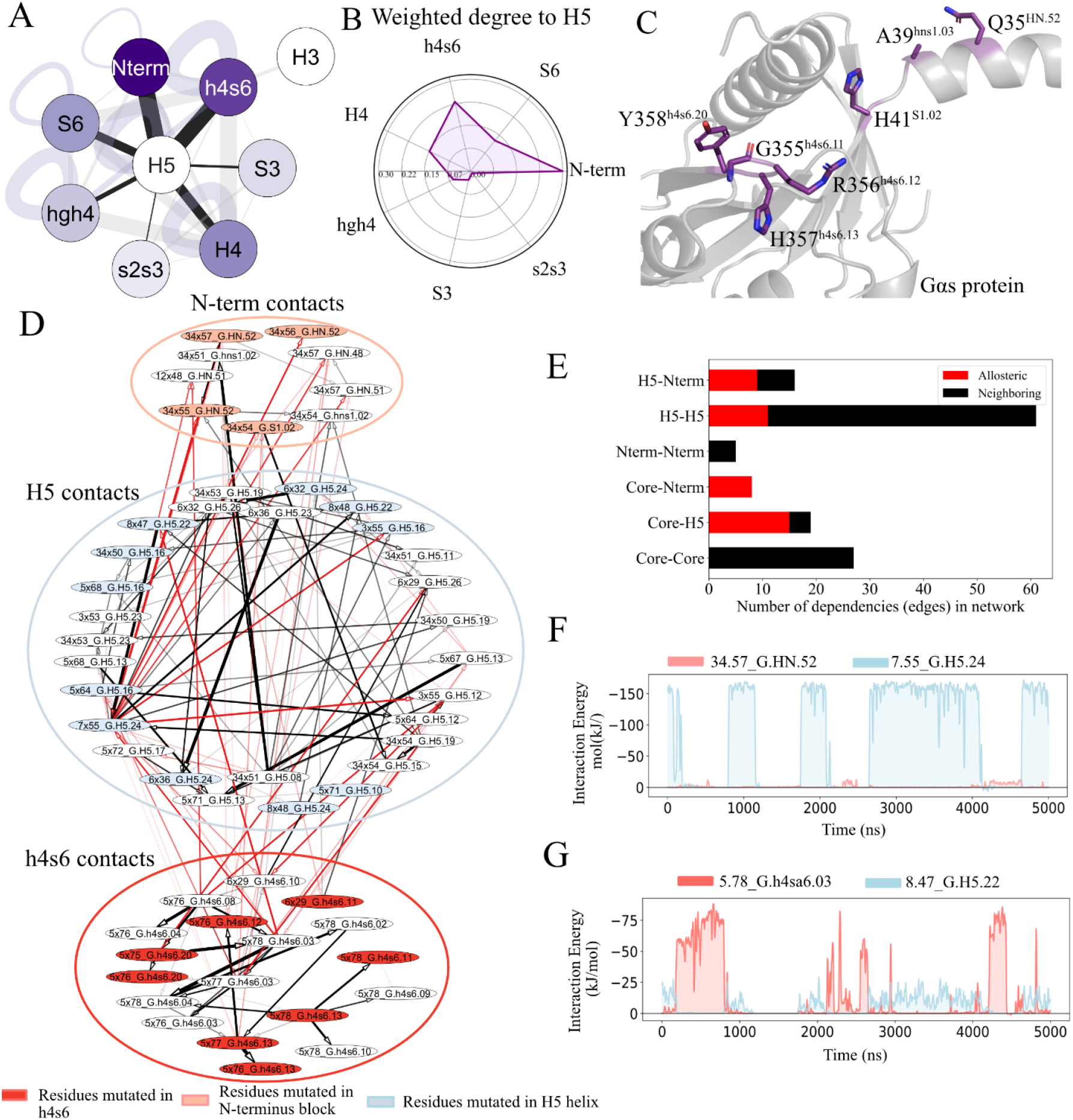
**A.** Network representation of G protein structural regions and their cooperative interactions with the H5 helix. Nodes represent distinct structural regions of the G protein, with connections (edges) indicating cooperative interactions with H5-helix contacts. Arrow thickness corresponds to the weight of cooperativity, with larger arrows indicating a stronger influence on the H5 helix. **B**. Radial plot of weighted degree of each structural region of G protein to H5 helix. **C**. Structural representation of core regions selected for mutation, including the N terminus (HN helix and hns1 loop) and the h4s6 loop. Mutated residues are colored purple and represented as sticks. **D**. Excerpt from subnetwork of cooperative interactions of β_2_AR-Gs protein system focusing on N-terminus, h4s6, and H5 helix regions. Arrows’ width is proportional to edge strength, color represents dependency type. Nodes containing mutated G protein residues are colored in: light blue – H5 helix residues, red – h4s6 residues, maroon – N-terminus. **E**. Number of allosteric and neighboring dependencies between H5, h4s6, and N-terminus regions in β_2_AR-Gs protein subnetwork of cooperative interactions depicted in G. Allosteric dependencies are red, neighboring – black. **F**. Interaction energy (kJ/mol) of N-terminus contact 34.57_G.HN.52 (maroon) and H5 contacts 7.55_G.H5.24 (blue) represented as time series. G. Interaction energy (kJ/mol) of core contact 5.78_G.h4s6.03 (red) and H5 contacts 8.47_G.H5.22 (blue) represented as time series.

To quantify the allosteric dependencies between the H5 helix and various core structural regions of the Gα subunit, we first categorized each node in the BN model by the Gα structural segment it involves (for example, “H5”, “H4”, “hgh4” nodes). We then identified all edges in the network that connect one node classified as H5 with another node from a core region. The sum of these edge weights serves as a measure of how strongly the core region allosterically affects the H5 helix and vice versa (Fig. 3A). This score quantifies how strongly each core region influences the H5 helix; a higher value indicates a greater cooperative impact on the H5 helix and, consequently, a more significant role in selective Gα protein coupling. Our findings revealed that both the h4s6 and N-terminus regions exert a strong cooperative influence on the H5 helix (Figure 3B), which guided our selection of specific regions for mutational analysis aimed at functional switching.

Based on this analysis, we performed mutations targeting the N-terminus and h4s6 sections of the core mutant (Table 1, Figure 3C) and tested each mutant’s ability to couple to β_2_AR and V_1A_R. The results in Fig. 2F show reduced isoproterenol-induced cAMP accumulation for both mutants; results in Fig. 2H indicate an increase of AVP-induced V_1A_R cAMP accumulation through Gαs N-terminus, h4s6, and N-terminus+h4s6, and the full-core-containing mutants. It is also observed that the combination N-terminus+h4s6 exhibits more significant cAMP accumulation than the Gαq-like H5 mutations, indicating that cooperative interactions within the core region may play a crucial role in modulating coupling selectivity. This observation aligns with our previous findings that the core region is a major site of cooperativity in both Gαs and Gαq proteins and suggests that targeting this region can significantly impact coupling dynamics.

To understand the mechanism behind these functional changes, we analyzed the dependencies between the mutated regions and the H5 helix. We enriched our BN graph with structural information, categorizing dependencies as either neighboring (distance between contacts is less than 10Å) or allosteric (>10Å). Our analysis revealed that the Gαs protein exhibits numerous allosteric dependencies between the core/N-terminus and H5 helix, indicating that changes in one region can propagate to influence another (Figure 3D, E). By focusing on the interaction subnetwork of regions where block mutations were performed (Figure 3E), we found that many of the mutated residues either directly connect to H5 helix contacts or are in the close probabilistic neighborhood.

To further demonstrate how these contacts allosterically influence each other during MD simulations, we calculated interaction energies for key contact pairs: 34.57_G.HN.52 - 7.55_G.H5.24 (N-terminus-H5 helix pair, Fig. 3F) and 5.78_G.h4s6.03 - 8.47_G.H5.22 (core-H5 helix pair, Fig. 3G). For the N-terminus-H5 helix pair, there is a direct dependency reflected in their interaction energies: when the interaction energy of the H5 contact is high, the energy of the N-terminus contact is low, and vice versa. This negative correlation suggests that one interaction can sustain coupling with the receptor. When we introduced a Gαs to Gαq switching mutation at HN.52, it enabled the Gαq-like Gαs protein to adopt a conformation conducive to successful Gαq-coupled protein interactions. For the core-H5 helix pair, although G.h4s6.03 could not be directly mutated due to its absence in Gαq proteins, its parent contact (5.75_G.h4s6.20) contains a mutated residue that encodes an undirected correlation between 8.47_G.H5.22 and 5.75_G.h4s6.20 via 5.78_G.h4s6.03. Consequently, a mutation in G.h4s6.20 indirectly altered the state of 5.78_G.h4s6.03, ultimately impacting H5 helix contacts.

Together, these analyses demonstrate that cooperative dependencies among core regions, N-terminus, and H5 helix are critical for mediating functional switching in G protein coupling to GPCR. By introducing targeted mutations in these cooperative regions, we have shown how specific residues can influence interdependent interactions dynamically, leading to altered coupling profiles. These findings further underscore the importance of understanding both local and allosteric dependencies for effectively modulating GPCR-Gα protein interactions.

### Dynamics of Gαs core-Gαq H5 chimera in comparison to wild type Gαs and Gαq proteins

Gα chimeras have been used extensively to stabilize and solve three-dimensional structures of GPCR:G protein complexes (35, 36). To understand the dynamics of the Gαq H5 chimera compared to the respective wild-type (WT) Gα proteins, we performed MD simulations and structural analyses on GPCR:Gα chimeras as well as GPCR:Gα wild type complexes. While the amino acid sequences in the core of Gαs, Gαq and Gαi are ∼55% similar, their three-dimensional structures show significant differences (Fig. S3A). To understand the differences in the dynamic ensemble, we performed 5 µs MD simulations for the serotonin 5HT_2A_ receptor complexed with the chimera Gαs core-Gαq H5. The results from these simulations were compared to β_2_AR-Gαs WT and H_1_R-Gαq WT complexes to assess the influence of the core on H5 helix dynamics. We clustered the conformational ensemble for each system based on the coordinates of the backbone atoms of the Gα protein. Representative structures extracted from the most populated clusters were analyzed to assess conformational similarities and differences (Fig. 4A-D).

**Figure 4.**
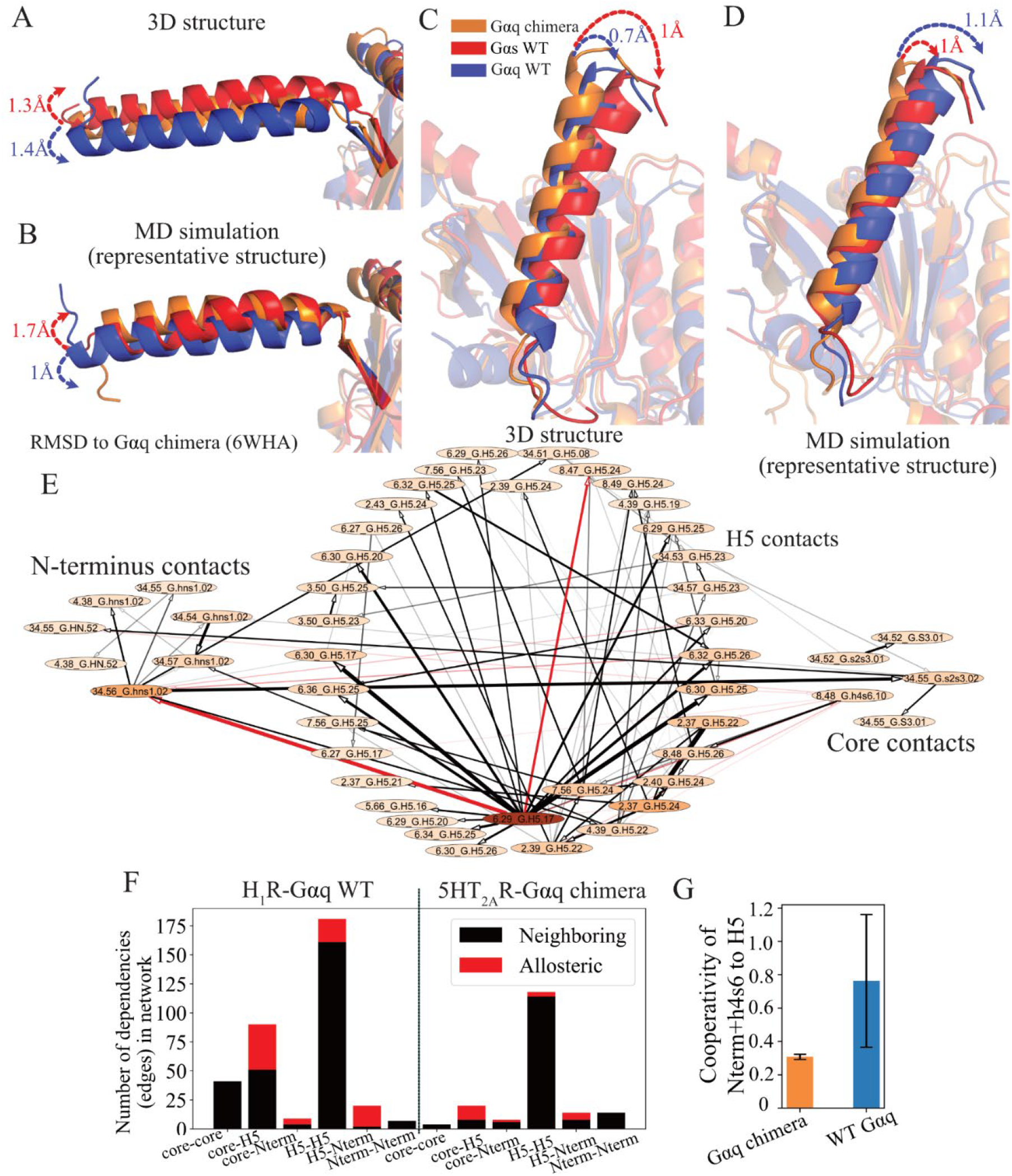
Dynamics and cooperativity in Gα chimera compared to the wild type Gα. **A.** Comparison of the N-terminus position in the three-dimensional structures of Gαq chimera (Gαs core-Gαq H5) from serotonin 5HT_2A_ (pdb id: 6WHA (orange)), Gαs WT from β_2_AR coupled to Gαs (pdb id:3SN6 (red)), and Gαq WT from histamine H_1_ receptor (pdb id:7DFL (blue)). Proteins are aligned by the backbone. **B**. Comparison of representative structures from MD simulations of Gαq chimera from 6WHA (orange), Gαs WT from 3SN6 (red), and Gαq WT from 7DFL (blue) proteins. N-terminus region is highlighted (opaque cartoon). **C**. Comparison of the H5 helix in the Gαq chimera from 6WHA (orange), Gαs WT from 3SN6 (red), and Gαq WT from 7DFL (blue) proteins. Proteins are aligned by the backbone. **D**. Comparison of representative structures from MD simulations of Gαq chimera from 6WHA (orange), Gαs WT from 3SN6 (red), and Gαq WT from 7DFL (blue) proteins. H5 helix is highlighted (opaque cartoon). **E**. Bayesian Network graph of cooperative interactions in the 5HT_2A_-Gαq chimera. Allosteric dependencies are shown as red arrows, neighboring – black arrows. Nodes are colored proportionally to a weighted degree. **F**. Bar plot showing the number of allosteric and neighboring interactions between core, N-terminus and H5 helix in H_1_-Gαq WT and 5HT_2A_-chimeric Gαq subnetwork of cooperative contacts. **G**. Bar plot showing the total cooperativity score between cooperative contacts in Nterm+h4s6 structural regions and H5 helix. Blue color corresponds to networks of Gαq WT: H_1_R and AT_1_R, orange color corresponds to chimeric Gαq proteins: 5HT_2A_ and M_1_R.

Comparison of the three-dimensional structures of these three receptors showed that the N-terminus of the chimera was positioned closer to the Gαs WT protein (Fig. 4A). Over the course of the MD simulation, the N-terminus shifted to a conformation resembling that of the Gαq WT protein. This transition was marked by a reduction in root-mean-square deviation (RMSD to PDB ID: 6WHA) from 1.4 Å to 1 Å (Fig. 4B). This transition highlights the dynamic flexibility of the chimera and its ability to adopt a Gαq-like state.

The H5 helix of the chimera closely aligned with the Gαq WT protein in the three-dimensional structure with an RMSD of less than 1 Å (Fig. 4C). However, during the simulation, the H5 helix diverged slightly from the Gαq WT conformation while maintaining a stable RMSD relative to the Gαs WT protein (Fig. 4D). This divergence likely stems from interactions between the base of the H5 helix and the core, where the chimera retained Gαs-like features. Such interactions likely influence the hinge region between the H5 helix and the core, as evidenced by residue contact patterns observed during the simulations (Fig. S3B).

We performed BN analysis of the residue contacts in 5HT_2A_-Gαq chimera interface to investigate its cooperativity and interdependencies, following methods described for the WT systems. The resulting cooperative interactions in the chimera are predominantly in the H5-mediated contacts, with minimal contributions from the core and N-terminus, in contrast to those in the WT Gαq or Gαs coupled complexes (Fig. 4F). Moreover, the number of allosteric dependencies between core and H5 helix was reduced in chimera BN compared to Gαq WT, which might indicate the loss of cooperative cross-talk between these two regions. To further support this observation, we measured the total cooperativity score of N-terminus and h4s6 regions to H5 helix in Gαq WT (H_1_R:Gαq protein, AT_1_R:Gαq protein, Fig. S4A,B) and chimeric Gαq proteins (5HT_2A_-miniGq, M_1_R-Gαq-Gαi1 chimera, Fig. S4C,D). As evident in Fig. 4G, chimeric Gαq protein shows lower cooperativity between Nterm+h4s6 and H5 helix than wild-type Gαq proteins. This further demonstrates that the chimera shows a decreased allosteric influence among Gα structural regions that can possibly lead to differences in Gα coupling to GPCRs. These findings align with the previous findings that the H5 helix is the most crucial region in Gαq proteins, compared to Gαs proteins (4, 13). In a prior BRET-based study (4), the authors tested all pairwise chimeras (i.e., swapping the helix 5 tip among Gαs, Gαi, Gαq, Gα12) for 12 GPCRs (4 Gi-, 4 Gs-, 4 Gq-, and 2 G12/13-coupled). They found that most receptors require both the Gα core and the H5 tip for efficient coupling. Notably, two Gαq-coupled receptors (H_1_R and endothelin A receptor) showed equal activity with either Gαq or Gαs cores, whereas Gαs-coupled receptors had significantly reduced efficiency with a non-cognate core. Overall, these findings suggest Gαq coupling depends more on the H5, whereas Gαs coupling is largely governed by the Gα core.

Our analyses demonstrate that the structural and dynamic integrity of the H5 helix is essential for Gαq-like signaling. While the core influences interactions at the base of the H5 helix to be more Gαs-like, it does not significantly alter the GPCR-G protein coupling interface. Critical H5 contacts responsible for Gαq-specific signaling are retained in the chimera. These results align with experimental evidence showing that Gαq proteins are less affected by modifications in the core (4), and underscore the pivotal role of the H5 helix in Gαq coupling specificity, highlighting its resilience to structural perturbations in the core (4).

## Discussion

Selectivity in protein-protein interactions (PPIs) is regarded as a property determined by the structure and dynamics of residue contacts at the interface. In contrast, protein allostery is appreciated as the communication between distal interaction sites that can influence PPIs. Here, we use a novel Interpretable Bayesian Network Model to highlight the pivotal role of multi-residue cooperativity, especially involving distal residue blocks in shaping the coupling selectivity. Using the thoroughly investigated GPCR-G protein interface as a model PPI that represents a superfamily of 800 receptors and 20 distinct Gα proteins, we reveal the complex interplay of allostery within the G protein in determining coupling selectivity. The BN model and the allostery concepts presented here are generalizable to any PPI.

The results from our MD simulations and BN modeling provide compelling evidence that cooperative contacts within the GPCR:Gα protein interface are essential for maintaining the stability and functionality of these complexes. Notably, while a substantial proportion of contacts are shared across different Gα protein subtypes, it is the subtype-specific interactions that predominantly drive cooperativity. This observation aligns with previous studies highlighting the importance of specific amino acid residues in mediating selective interactions, particularly in regions such as the H5 helix and various intracellular loops (4–7, 37).

Our analysis reveals that the distribution of cooperative contacts varies significantly among different G protein subtypes. For instance, Gαi and Gαq coupled receptors exhibit high cooperativity contacts between the H5 helix of the Gα protein and specific transmembrane domains of GPCRs, while Gαs coupled receptors demonstrate a more uniform distribution of cooperative interactions among H5 and core regions. These differences in the location of the cooperativity contacts suggest that specific structural features are tailored to facilitate selective signaling pathways, which is crucial for understanding how GPCRs can elicit distinct physiological responses despite sharing overlapping structural characteristics.

Although the Gα core has been shown to be important for coupling selectivity (4), our mutational analysis provides insights into how targeted modifications in the Gα core regions can influence Gα coupling profiles. By engineering mutations based on cooperativity hotspots, we successfully altered the coupling behavior of Gαs to mimic that of Gαq (Figure 2). This ability to switch coupling preferences through predicted mutations emphasizes the potential (i) of interpretable network-centric machine learning, specifically BN models, for identifying cooperative interactions in any protein-protein interface (ii) for developing therapeutics that can selectively modulate GPCR signaling pathways by targeting cooperative sites within the G protein structure and (iii) for delineating the allosteric dependencies between various structural determinants in the Gα core to the tip. These findings give us the ability to predict block mutations that show co-dependencies, a concept that will be useful in protein design. Thus, using the GPCR:G protein complexes as a model we have uncovered the cooperativity and allosteric dependencies about distant structural regions in Gα subtypes, a concept that is applicable to other protein interfaces.

The application of BN modeling has proven invaluable in elucidating the complex interdependencies among residue pairs at the GPCR:Gα protein interface. By representing these interactions as direct probabilistic dependencies within a graphical model, we have been able to distinguish true cooperative interactions from spurious correlations that often arise in conventional analyses. This approach not only enhances our understanding of how local and long-range interactions contribute to signaling specificity but also provides a framework for identifying critical residues involved in allosteric communication within any protein complex.

In summary, this study sheds light on the multifaceted nature of GPCR:Gα protein coupling dynamics, revealing that cooperative interactions are fundamental to achieving selective signaling outcomes. By integrating MD simulations with BN modeling, we have bridged the gap between static structural data and dynamic functional insights, paving the way for future research aimed at understanding and manipulating GPCR signaling pathways. Our results suggest that targeting cooperative sites within G proteins could represent a promising strategy for designing novel therapeutics with enhanced specificity and efficacy in modulating cellular responses. Future studies should explore additional factors such as ligands, allosteric modulators, and post-translational modifications influence this cooperativity and can investigate how these dynamics can be harnessed to develop more effective pharmacological interventions.

## Materials and Methods

### Preparation of GPCR:G protein complexes for MD simulations

We used experimental three-dimensional structures of the GPCR:G protein complexes as starting structures for the MD simulations: 6GDG (27), 6D9H (30), 6G79 (31), 7F6G (29), 7DFL (28), 3SN6 (26), 6WHA (36), 6OIJ (35). Mutations in both receptors and Gα subunits found in the three-dimensional structures were reverted to their wild-type sequences using Maestro (Schrödinger Release 2020-1). Mutations in the chimeric Gαq proteins from 6WHA and 6OIJ were kept unchanged. GDP was missing in all complexes simulated. Ligands were parameterized using ParaChem (https://cgenff.umaryland.edu, (38)). Missing sidechains and loops with fewer than five residues were added to the GPCR:G protein complex structures. Localized minimization of residues within 5 Å of mutation sites was carried out using MacroModel, with backbone atoms positionally restrained. Chain termini were capped with neutral acetyl and methylamide groups, and histidine protonation states were assigned using Maestro’s protein preparation wizard. The prepared complexes were embedded in an explicit POPC bilayer membrane using the PPM 2.0 module from the Orientation of Proteins in Membranes (OPM) tool (39). The systems were then solvated with TIP3P water containing 0.15 M NaCl using CHARMM-GUI (40–42). The final system dimensions were approximately 120 Å × 120 Å × 170 Å. All components were parameterized with the CHARMM36m force field (43).

### MD simulations for GPCR:G protein complexes

MD simulations were performed using GROMACS2019 (44, 45) with a 2 fs integration time step. Systems were initially minimized under positional restraints of 10 kcal/mol·Å^2^ applied to heavy atoms of the GPCR, Gα subunit, ligand, and lipid molecules. Following minimization, the systems were heated from 0 K to 310 K over 1 ns using the NVT ensemble and the Nosé-Hoover thermostat. Equilibration was then conducted under NPT conditions, beginning with positional restraints of 10 kcal/mol·Å^2^ that were progressively reduced to 1 kcal/mol·Å^2^ in 1 kcal/mol·Å^2^ increments, with each increment simulated for 5 ns. A final unrestrained equilibration run of 50 ns was performed, and the last snapshot from this stage was used to initialize five independent production simulations of 1 μs each, starting with different randomized velocities. Pressure coupling was maintained at 1 bar using the Parrinello-Rahman barostat (46), and nonbonded interactions were truncated at 12 Å. Long-range van der Waals interactions were computed using the Particle Mesh Ewald (PME) method (47). The LINCS algorithm was applied to constrain all bonds, and the system stability was monitored by tracking the root-mean-square deviation (RMSD) of receptor and Gα protein backbone atoms over time (Fig. S1A). 5 production simulations were concatenated together with a snapshot frequency of 200ps for the final analysis. RMSD for TM backbone atoms in the transmembrane helices demonstrating the convergence of MD simulations is shown in Fig. S1A.

### RMSD-based clustering of conformation ensembles

To obtain representative snapshots from MD simulations demonstrated in Fig.4A-D, we used gmx cluster to calculate the most populated cluster representatives for G protein conformations at 3Å RMSD cutoffs. Complexes in MD simulations were aligned to the backbone of helix regions of Gα protein from the starting structure using rot+trans fitting.

### Analysis of pairwise contacts between GPCR and Gα subunits

The Python-based tool “GetContacts” (https://www.github.com/getcontacts, (32)) was used to identify intermolecular contacts between GPCR and Gα subunits. Contact types analyzed included salt bridges (<4.0 Å between anion and cation), hydrogen bonds (<3.5 Å between donor and acceptor, angle <70°), van der Waals interactions (<2 Å distance), pi-stacking interactions (<7.0 Å between aromatic centers, angle <30°), and cation-pi interactions (<6.0 Å between a cation and an aromatic ring centroid, angle <60°). All interaction types were analyzed across the full 5 μs trajectories, excluding water and ions. Residues in the GPCR and Gα subunits were matched to their respective domains using generic numbering systems (Ballesteros-Weinstein for GPCRs and Common G protein numbering for Gα). A one-hot encoding approach was applied to generate binary interaction fingerprints for each frame, where “1” indicated the presence of a contact and “0” its absence.

### Bayesian Network Analysis of interactions

Binary interaction fingerprints were analyzed using BaNDyT (https://github.com/bandyt-group/bandyt) (33), a Bayesian network (BN) modeling tool for MD simulation trajectories. Independent BN models were generated for each GPCR:G protein complex using a search-and-score approach. The Minimum Uncertainty (MU) scoring function was employed, with 50 random restarts to ensure convergence (48). Nodes in the BN corresponded to residue pairs, while edges represented their direct probabilistic dependencies, with edge strengths quantified by MU on a commensurate bounded scale. The weighted degree, calculated as the sum of edge strangths for each node, was used to rank contact pairs by cooperativity. Top-ranked contacts (top 25%) were compared across different Gα subtypes. Networks were visualized using Cytoscape 3.9.1 (49).

### Robustness assessment of Bayesian Networks

The robustness of the BN models was evaluated using randomization and sensitivity analyses. For randomization, 1,000 perturbations were introduced into the network by randomly adding or removing edges, and the MU scores of perturbed models were compared to the original (48). Sensitivity testing involved resampling trajectory frames 1,000 times with replacement and recalculating model scores against the original network topology. Results were evaluated for consistency within the 0.99 confidence interval.

### Calculation of residue interaction energy

Interaction energies between residue pairs were analyzed using ensemble trajectories. The GROMACS “energy” (44, 50) tool was utilized to compute the total nonbonded interaction energy, which includes short-range coulombic and van der Waals forces within a 12 Å cutoff. These values were extracted from the energy log file and summed to determine the overall nonbonded interaction energy for each residue pair.

### Reagents

(−)-Isoproterenol (+)-bitartrate salt, X-tremeGENE HP DNA transfection reagent, forskolin FSK, and 3-Isobutyl-1-methylxanthine (IBMX) were purchased from Sigma Aldrich. Synthetic Arginine 8 Vasopressin (AVP) was purchased from GenScript. Cell lines: HEK293-Flp-in (Invitro-gen) and HEK293ΔGsix, HEK293 absent of six functional G proteins (ΔGs, ΔGolf, ΔGq/11, ΔG12/13) (51). Human V1AR and G protein constructs were cloned into pcDNA5/FRT backbone following standard cloning procedures. The V1AR SPASM sensor constructs were assembled with G-S-G repeats in between domains (Receptor-4xGSG-mCetrine-4xGSG-10nm ER/K linker-4xGSG-mCerulean-4xGSG-G protein). The constructs expressing the alpha subunit of G protein were ensembled as mCerulean-4xGSG-G protein.

### Cell culture

HEK293-Flp-in (Invitro-gen) and HEK293ΔGsix were cultured with DMEM containing 10% FBS, 4.5 g/liter D-glucose, 1% Glutamax, 20mM HEPES, pH 7.5, at 37 °C in a humidified atmosphere at 5% CO2. HEK293ΔGsix cells culture media was supplemented with penicillin and streptomycin. Cells were used during passages 10-27 and seeded at 30% confluency in tissue culture-treated six-well plates. The cells were allowed to adhere for 16-20 hours before transiently transfecting them with X-tremeGENE HP DNA transfection reagent. The transfection conditions were optimized to 1 μg DNA, 3 μL transfecting reagent, and varying length of transfection (20-36 hours) to provide consistent expression levels for all constructs.

### cAMP accumulation assay

Cells were harvested post-transfection, and ligand-dependent cAMP accumulation was measured using the cAMP Glo assay (Promega). In brief, 1 mL of media was removed from the well to be harvested and gently resuspended with the remaining volume. Harvested cells were centrifuged at 300g for 3 min, and media was removed by vacuum manifold. The cell pellet was resuspended in 1 mL of cAMP assay buffer (PBS with 0.5mM ascorbic acid, 0.2% (w/v) glucose). The cells were washed once by repetition of this procedure. Cell density was measured with Countess II and diluted to 3.0×106 cells per mL. Expression of the selected G protein or V1R tethered G protein was measured through the fluorescence spectra of mCerulean fused in these constructs. The metric used to assess the expression was the ratio between the mCerulean peak emission (475 nm with 430 nm excitation) and optical density emission (450 nm with 430 nm excitation)

Resuspended cells were incubated in an opaque 384-well flat-bottom plate (Greiner Bio-One) with an equal volume of 2x concentration of ligand for a final concentration of 10 μM Isoproterenol or 100 nM AVP, supplemented with 0.5 mM IBMX to inhibit phosphodiesterase activity. After 15 minutes at room temperature of incubation, cells were lysed and processed for the cAMP-Glo Assay (Promega) following the manufacturer’s instructions. Luminescence was measured on a Tecan Spark plate reader (500 ms integration, one measurement per well). Each condition was performed on four independent wells, and the experiment was performed at least 3 times (n >3). Data was illustrated and analyzed using GraphPad Prism 10 (version 10.4.0).

## Acknowledgments

We would like to thank Asuka Inoue for providing HEK293ΔGsix cells culture. This work was supported in part through the City of Hope’s High Performance Computing resources, services, and staff expertise provided under the ETG. This work was funded by grants from the National Institutes of Health R01-GM117923 to N. V. and S.S., R01-LM013876 to N. V., A. R., S.B., and R01-LM013138 to A. R. The content is solely the responsibility of the authors and does not necessarily represent the official views of the National Institutes of Health. Additional support is acknowledged by Dr Susumu Ohno Chair in Theoretical Biology (held by A.R.).

## Author Contributions

Conceptualization: S.B, A.R., N.V.

Methodology: E.M., E.R., S.B, A.R., S.S., N.V.

Investigation: E.M., E.R.

Visualization: E.M., E.R.

Formal analysis: E.M., E.R.

Funding acquisition: S.B, A.R., N.V.

Supervision: S.S., N.V.

Writing – original draft: E.M, E.R., N.V

Writing – review & editing: E.M, N.V.

## Data and materials availability

All data is included as supplementary files.

## Supplementary Materials

**Figure S1.**
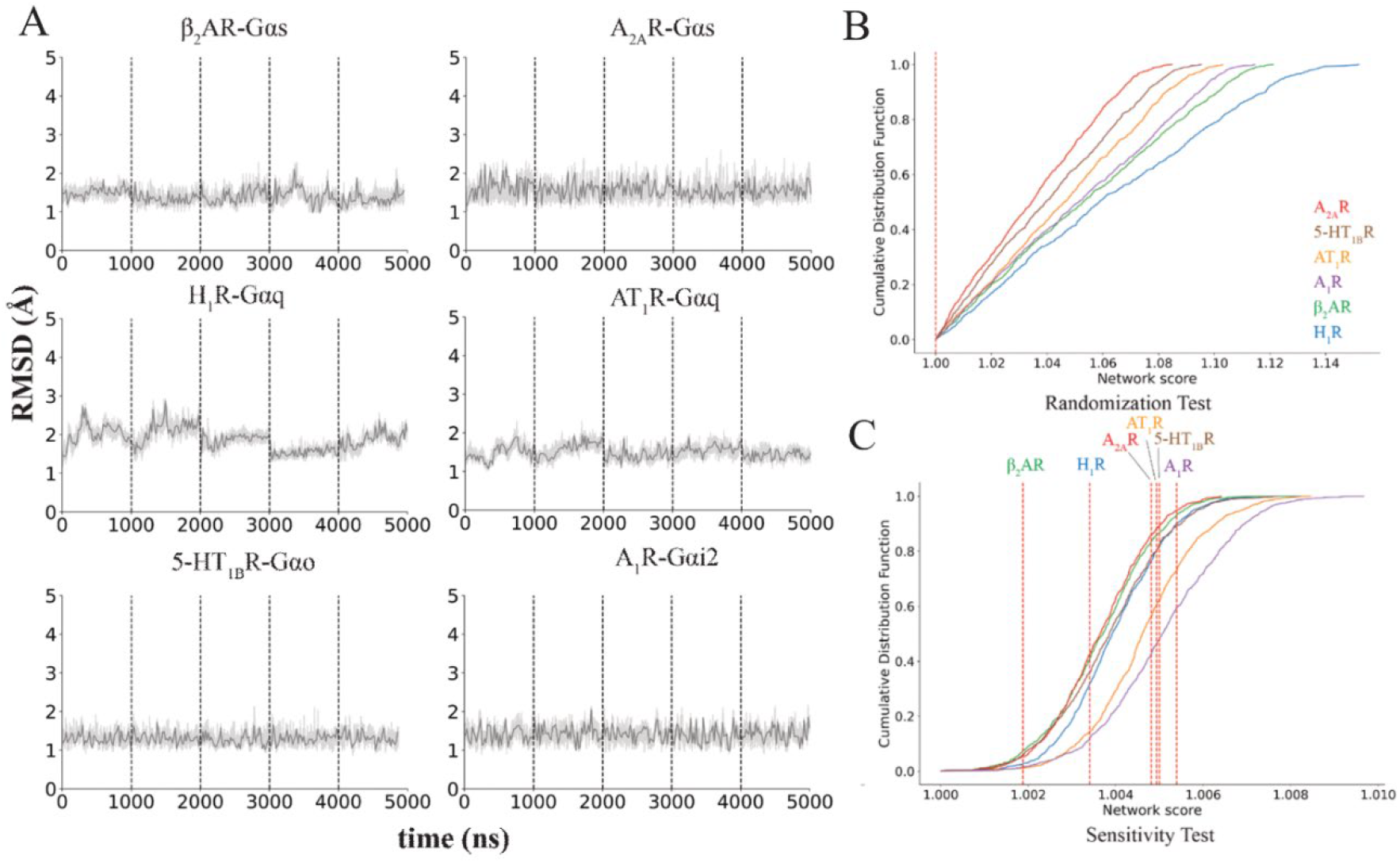
Analysis of GPCR:Gprotein interactions and cooperative contact properties. **A**. Root-mean-square deviation (RMSD) values in Å for TM backbone atoms in the transmembrane helices. **B**. Cumulative Distribution Function of all network scores with various topology perturbations. Red dashed line indicates the placement of score of the original networks. **C**. Cumulative Distribution Function of all network scores with data resampling. Red dashed line indicates the placement of score of the original networks.

**Figure S2.**
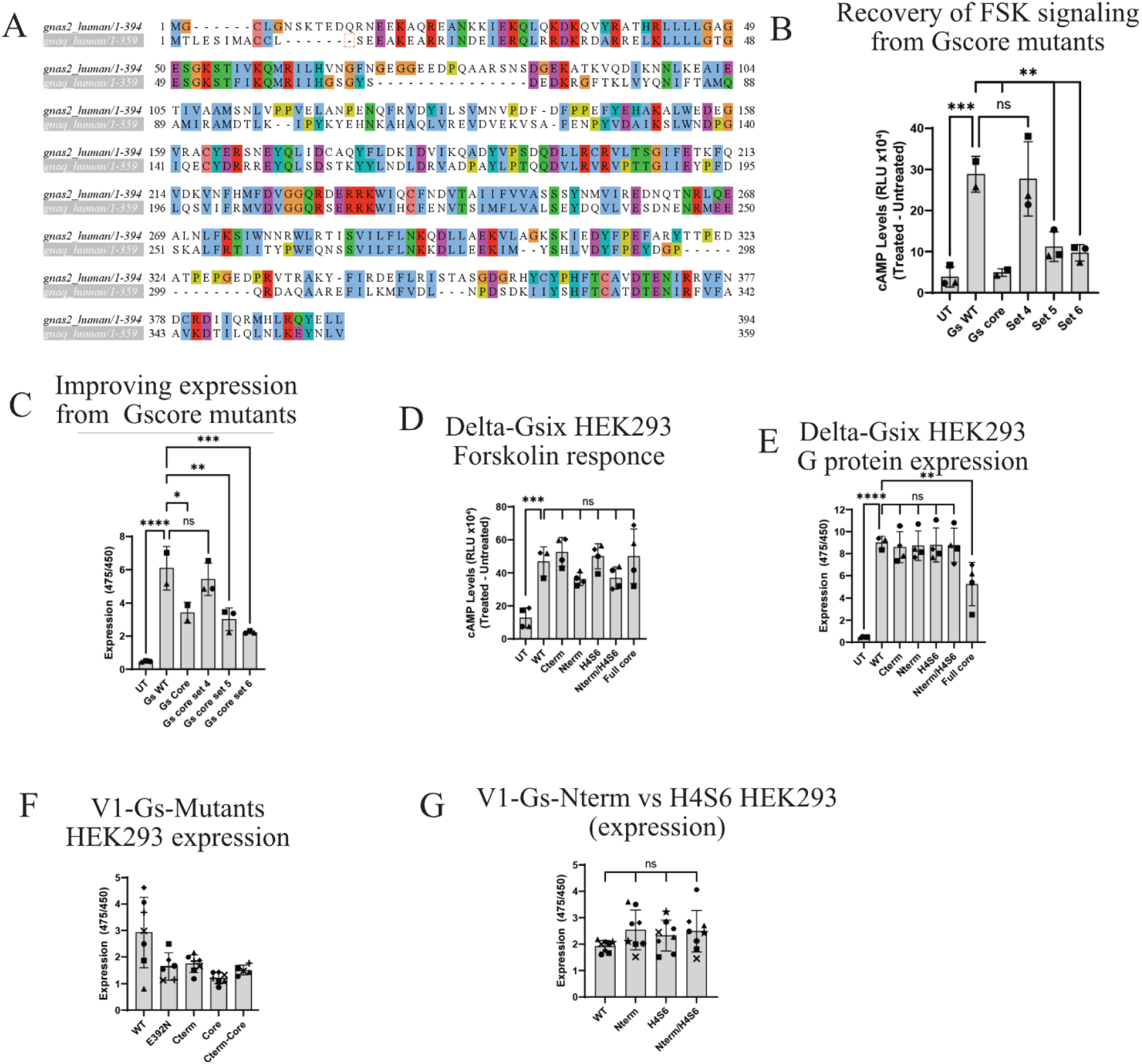
**A**. Multiple sequence alignment of Gs and Gq proteins built using Clustal Omega server. **B**. Forskolin signaling essay for wild type Gs protein and starting core mutant Gs protein. Set 4-6 correspond to subset of core mutations with following mutation blocks missing: set 4 does not contain H4 helix mutations, set 5 – N terminus, set 6 – h4s6 block. **C**. Expression of wild type Gs protein and starting core mutant Gs protein. Set 4-6 correspond to subset of core mutations with following mutation blocks missing: set 4 does not contain H4 helix mutations, set 5 – N terminus, set 6 – h4s6 block. **D**. Forskolin response of Gs protein mutants used in this study. **E**. Expression of Gs protein mutants used in this study. **F**. Expression of tethered V1R-Gs protein core and h5 helix mutants used in this study. **G**. Comparison of levels of expression of tethered V1R-Gs protein N terminus and h4s6 mutants.

**Figure S3.**
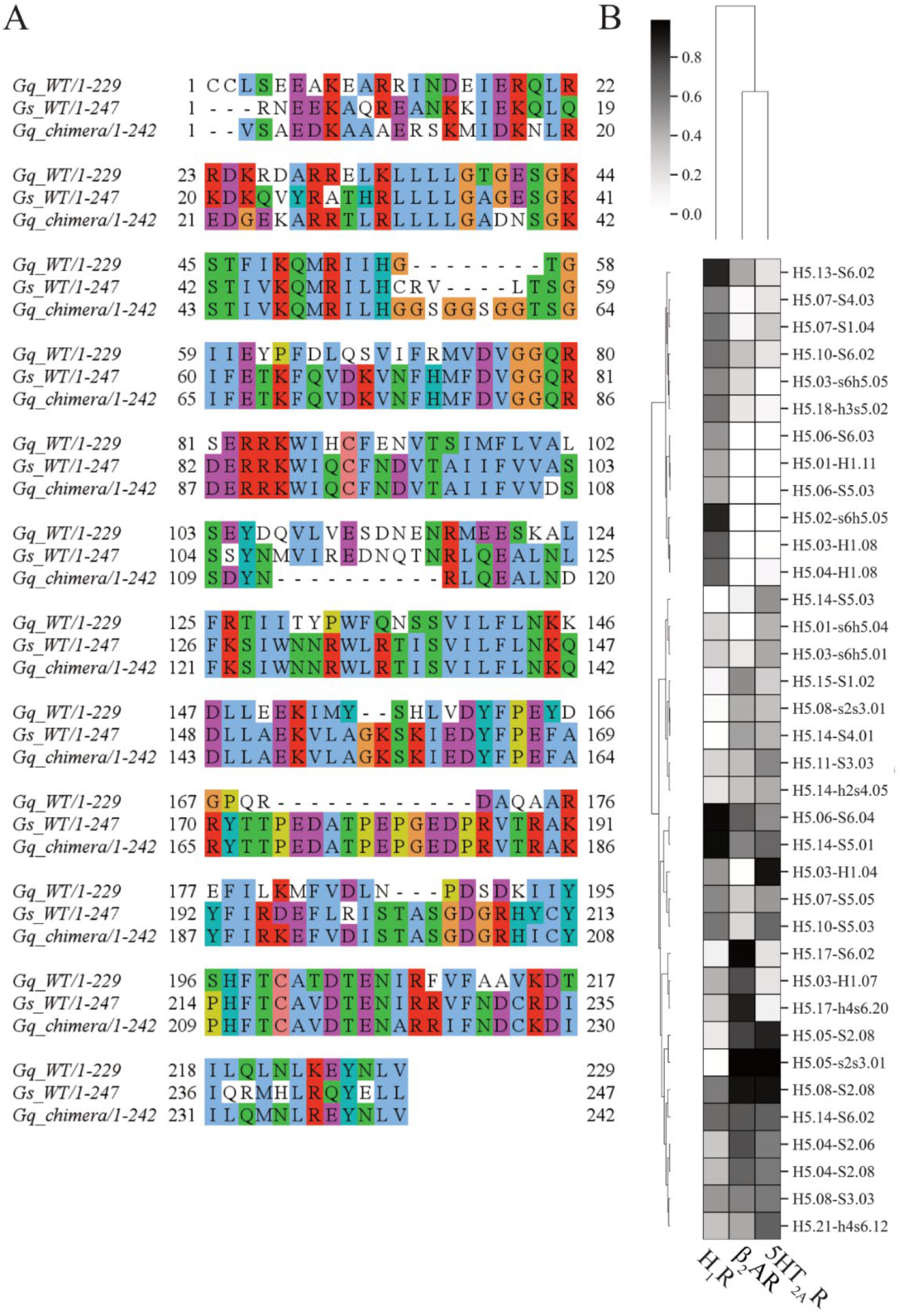
Structural and Cooperative Features of G Protein Core and H5 Regions. **A**. Multiple sequence alignment of Gs WT, Gq WT and Gs-core Gq chimera protein sequences built using Clustal Omega service. **B**. Contact persistency heatmap between the H5 helix and the core region in Gs WT, Gq WT and Gq chimera.

**Figure S4.**
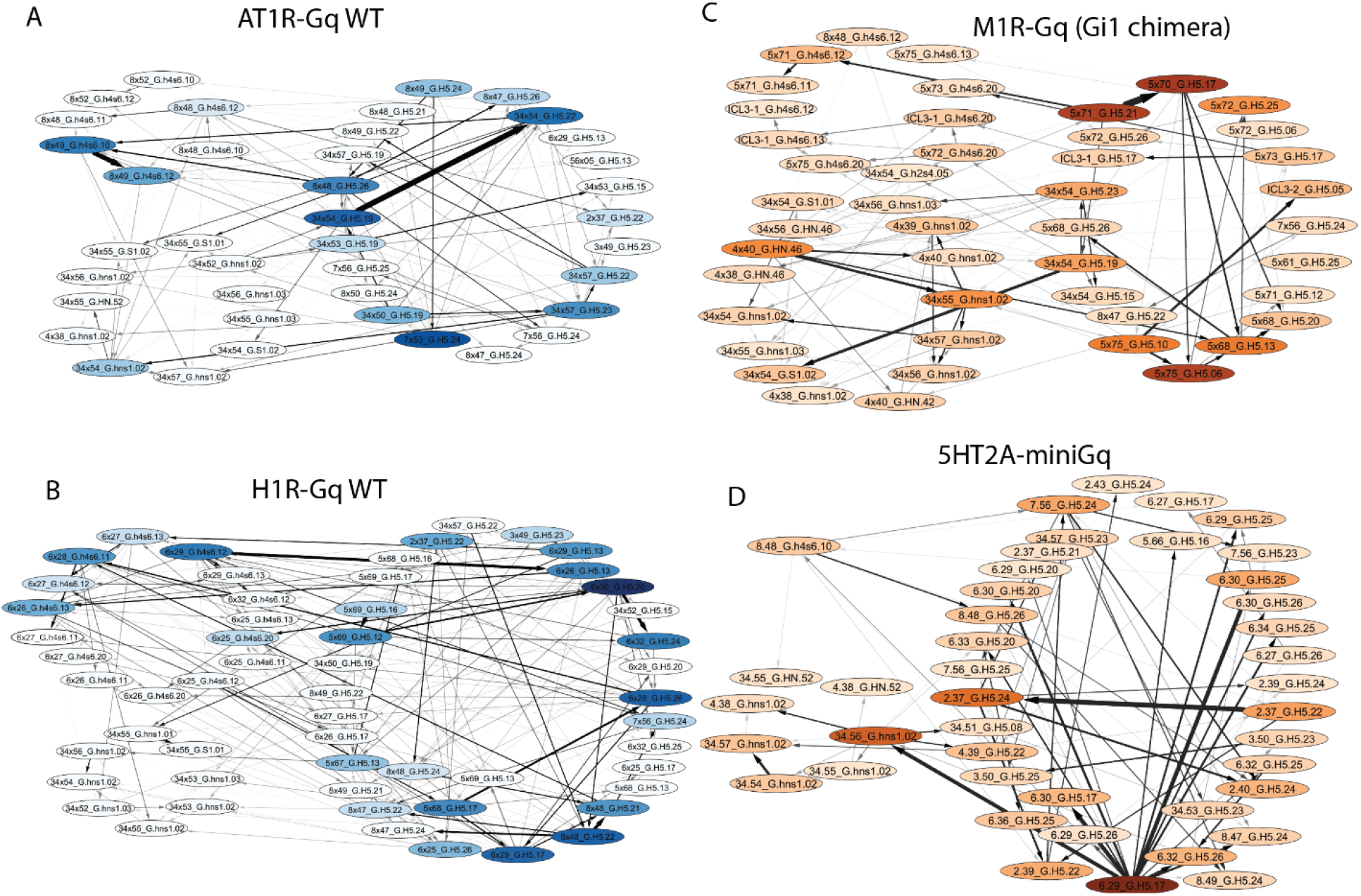
Subnetworks of N-terminus, h4s6 loop and H5 cooperative interactions of chimeric and wild type Gαq proteins. **A**. Subnetworks of N-terminus, h4s6 loop and H5 cooperative interactions of AT_1_R-WT Gαq protein. Color intensity is proportional to weighted degree, arrow thickness is proportional to edge weight **B**. Subnetworks of N-terminus, h4s6 loop and H5 cooperative interactions of H_1_-WT Gαq protein. Color intensity is proportional to weighted degree, arrow thickness is proportional to edge weight. **C**. Subnetworks of N-terminus, h4s6 loop and H5 cooperative interactions of M^1^R-Gαq-Gαi1 chimeric protein. Color intensity is proportional to weighted degree, arrow thickness is proportional to edge weight. **D**. Subnetworks of N-terminus, h4s6 loop and H5 cooperative interactions of 5HT_2A_-miniGq protein. Color intensity is proportional to weighted degree, arrow thickness is proportional to edge weight

